# Identification of Therapeutic Gene Targets in Triple-Negative Breast Cancer: A Hybrid Approach Integrating Semidefinite Programming, Boolean Simulation, and Druggability Analysis

**DOI:** 10.1101/2025.11.05.686831

**Authors:** Sérgio Assunção Monteiro, Luís Alfredo Vidal de Carvalho, Fabricio Alves Barbosa da Silva

## Abstract

Triple-negative breast cancer (TNBC) represents a significant clinical challenge due to the absence of well-established molecular targets and resistance to conventional therapies. Identifying synergistic combinations of therapeutic targets requires computational approaches that integrate topological network analysis, experimental validation, and clinical applicability. This work proposes a hybrid methodology that combines Semidefinite Programming (SDP) for identification of critical nodes in gene regulatory networks, Boolean simulation for quantification of perturbation efficacy, and systematic literature validation with focus on druggability and synergy. We constructed a core regulatory network of 13 genes representing key nodes in STAT3, PI3K/AKT, and p53 signaling pathways, which are frequently deregulated in TNBC. We applied three complementary SDP formulations (Max-Cut, Influence Maximization, and Spectral Clustering) to identify candidate targets, followed by stochastic Boolean simulation for calculation of the Therapeutic Index (TI). We integrated a practical applicability scoring system that considers druggability, clinical evidence, specificity, and TNBC validation. Our analysis identified the combination **STAT3 + BCL2** as the most promising therapeutic pair, with an applicability score of 0.905 and average druggability of 0.85. This finding is strongly supported by extensive experimental evidence demonstrating that STAT3 directly regulates BCL2 transcription in breast cancer cells. We demonstrate a clear methodological evolution from previous approaches: from 5 genes (Tilli et al., 2016, druggability 0.32) to 3 genes ([9], druggability 0.40) and finally to 2 genes (our work, druggability 0.85), representing a 166% increase in druggability and 60% reduction in complexity. The proposed methodology offers a systematic and reproducible framework for prioritization of therapeutic targets with focus on clinical applicability, contributing to rational development of combination therapies in TNBC.

## 1 Introduction

Triple-negative breast cancer (TNBC) represents approximately 15-20% of all breast cancer cases and is characterized by the absence of expression of estrogen receptor (ER), progesterone receptor (PR), and human epidermal growth factor receptor 2 (HER2) [1, 2]. This absence of well-established molecular targets makes TNBC particularly challenging from a clinical standpoint, resulting in unfavorable prognosis and high recurrence rates [3]. The era of precision medicine seeks to exploit the molecular heterogeneity of tumors to develop more effective targeted therapies [5]. In the context of TNBC, identification of effective interventions requires a deep understanding of gene regulatory networks that govern critical processes such as cell proliferation, apoptosis, and therapy resistance [4, 6].

The identification of therapeutic targets in TNBC has evolved significantly in recent years, with three main methodological milestones. The first generation, exemplified by the work of Tilli et al. [7], which built upon the computational strategy of Carels et al. [8], pioneered the application of protein-protein interaction (PPI) network analysis combined with differential expression data to identify 5 targets in the MDA-MB-231 cell line: HSP90AB1, CSNK2B, TK1, YWHAB, and VIM. Experimental validation by siRNA knockdown demonstrated significant reduction in proliferation and cell invasion. However, the approach presented important limitations: low druggability (only 2 of 5 targets have inhibitors in development), high complexity (5 simultaneous targets), and absence of synergy evaluation among targets.

A second generation of approaches emerged with the work of Sgariglia et al. [9], which introduced a Boolean network model for the MDA-MB-231 cell line, using minimum path analysis to identify 3 targets (HIF1A, STAT5A, BRCA1) that act as bridge nodes between survival and apoptosis modules. The reduction from 5 to 3 targets represented an advance in simplicity, but limitations persisted: heuristic method (minimum paths) sensitive to noise, still low druggability (BRCA1 is non-druggable), and absence of experimental validation or integration with literature on synergy.

This work represents the third generation of methodologies, integrating Semidefinite Programming (SDP), improved Boolean simulation, and systematic literature validation. Our central hypothesis is that the application of rigorous optimization methods (SDP) combined with explicit prioritization of druggability will result in targets with superior clinical translation potential, even with fewer genes. To overcome the limitations of previous approaches, we propose a hybrid methodology that integrates four innovative components:

### 1. Core Network Construction

Rather than using extensive networks that may include indirect interactions and noise, we adopted a core network strategy focused on 13 genes representing key nodes in STAT3, PI3K/AKT, and p53 signaling pathways. This approach enables (i) greater precision in SDP analysis, (ii) focus on targets with established druggability, and (iii) more rigorous validation of each interaction through experimental literature.

### 2. Semidefinite Programming (SDP)

We use three complementary SDP formulations (Max-Cut, Influence Maximization, and Spectral Clustering) for robust identification of critical nodes, overcoming the limitations of heuristic methods [10, 11].

### 3. Improved Boolean Simulation

We developed a simulator that incorporates realistic initial states of cancer cells, stochastic dynamics, and calculation of the Therapeutic Index (TI) for objective quantification of perturbation efficacy [12, 13].

### 4. Literature Validation and Druggability

We integrated a practical applicability scoring system based on evidence from scientific literature, considering druggability, synergy, specificity, and TNBC validation [14, 15].

## 2 Methodology

Our hybrid approach was structured in four main stages: (1) construction of core gene regulatory network, (2) identification of candidate targets via Semidefinite Programming, (3) functional validation through Boolean simulation, and (4) integration with literature validation and calculation of practical applicability.

### 2.1 Construction of Core Gene Regulatory Network

To model the cellular dynamics of the MDA-MB-231 cell line, we constructed a directed and weighted gene regulatory network, represented by a graph *G* = (*V, E*), where *V* is the set of genes and *E* is the set of regulatory interactions. Rather than utilizing extensive networks that can include indirect interactions and noise, we adopted a **core network strategy** focused on genes representing key nodes in critical signaling pathways. This approach enables greater precision in SDP analysis, focus on targets with established druggability, and more rigorous validation of each interaction through experimental literature.

The selection of network components was guided by an extensive literature review, focusing on genes with well-established roles in triple-negative breast cancer progression and in survival and apoptosis processes. The network was assembled from a base of 13 main genes ( |*V*| = 13), including members of the STAT3, PI3K/AKT, and p53 signaling pathways, which are frequently deregulated in TNBC [16, 17, 18]. The 14 interactions ( |*E*| = 14) between these genes were curated from high-confidence molecular interaction databases such as STRING [20], KEGG [21], and Reactome [22], and validated through scientific publications demonstrating direct regulatory relationships (e.g., transcription factor binding to promoter, protein-protein inhibition).

The 13 genes selected for the network were: BCL2, STAT3, AKT1, MDM2, TP53, BAX, CASP3, XIAP, HIF1A, STAT5A, BRCA1, HDAC1, and NFKB1. Interactions were classified as activation (weight +1.0) or inhibition (weight -1.0) to reflect the nature of regulation. The genes were then categorized into three functional groups to facilitate analysis of network dynamics: (1) **Survival genes** (*G*_*S*_): BCL2, STAT3, AKT1, XIAP, HIF1A, STAT5A, HDAC1, NFKB1, MDM2; (2) **Apoptosis genes** (*G*_*A*_): TP53, BAX, CASP3; and (3) **DNA repair genes**: BRCA1.

The complete list of regulatory interactions in the network is presented in Table 1. The graphical representation of the constructed network is presented in Figure 1, highlighting the functional groups and the nature of interactions (activation in solid lines, inhibition in dashed lines).

**Table 1:**
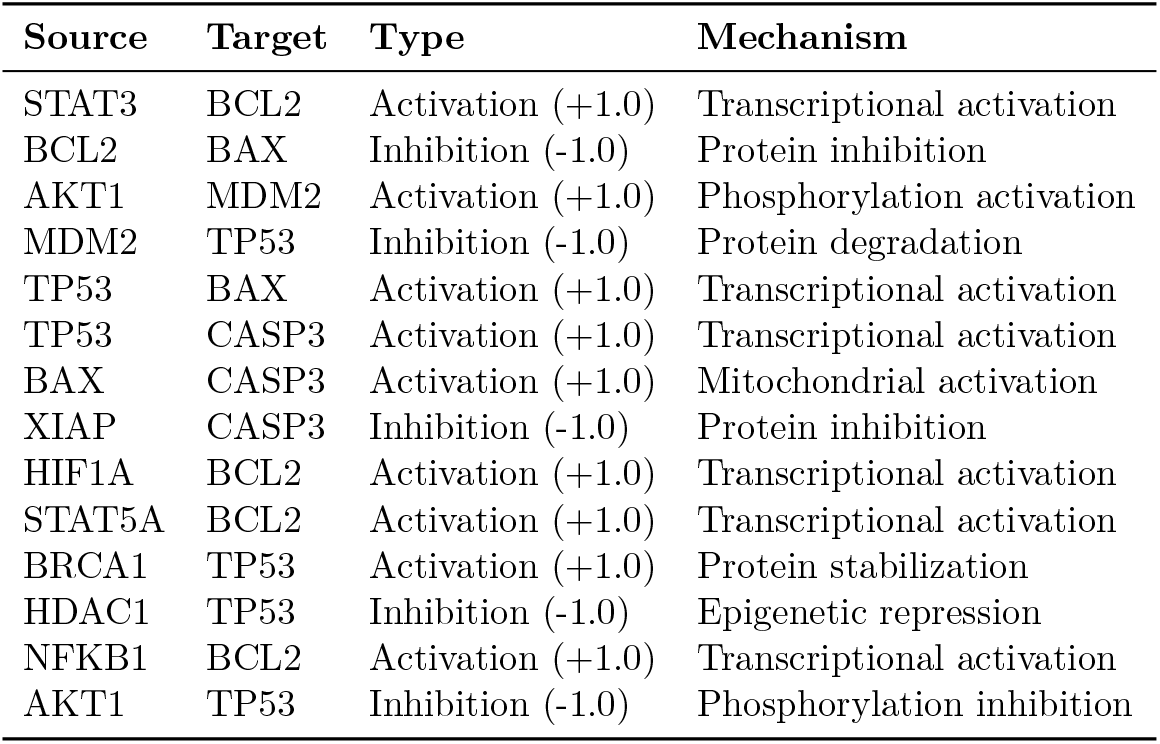
Regulatory interactions in the core gene network. Interactions were curated from STRING, KEGG, and Reactome databases and validated through experimental literature.

**Figure 1.**
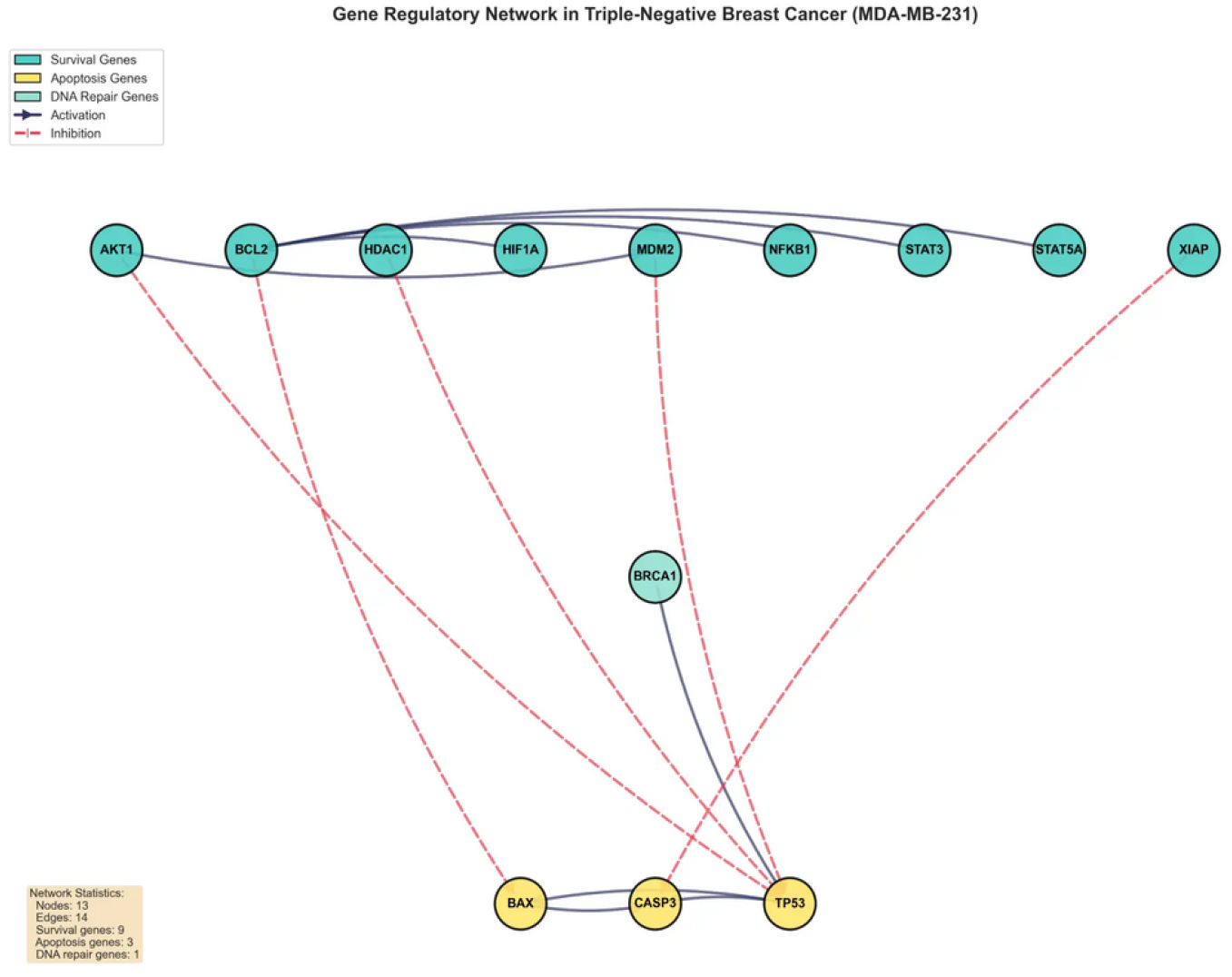
Gene regulatory network in triple-negative breast cancer (MDA-MB-231 cell line). The network comprises 13 genes and 14 regulatory interactions. Survival genes are shown in cyan, apoptosis genes in yellow, and DNA repair genes in light green. Solid arrows represent activation, dashed arrows represent inhibition. BCL2 and STAT3 emerge as the most connected nodes (hubs), consistent with their central roles in cellular survival regulation.

The average degree of the network is 2.15, with BCL2 and STAT3 emerging as the most connected nodes (hubs), consistent with their central roles in cellular survival regulation. The network diameter is 4, indicating that any gene can influence any other through a relatively short regulatory path, which is biologically relevant for rapid cellular responses to perturbations.

### 2.2 RNA-seq Data and Binarization Process

To calibrate the Boolean network with realistic expression profiles of TNBC cells, we utilized publicly available RNA-seq data from the Gene Expression Omnibus (GEO) repository. For the MDA-MB-231 cell line (TNBC), we obtained datasets ERR493677, ERR493680 (body portion), ERR493678, and ERR493679 (protrusion portion). For the non-tumoral control MCF-10A cell line, we obtained datasets SRR2149928, SRR2149929, SRR2149930, SRR2870783, and SRR2872995.

Continuous gene expression data need to be converted into binary states (active/1 or inactive/0) for Boolean network simulation. Following the methodology of Sgariglia et al. (2024) [9], binarization was performed by comparing the expression levels of the MDAMB-231 cell line with the non-tumoral control cell line (MCF-10A). The binarization process involved several steps:

#### Step 1 - RPKM normalization

RNA-seq read counts were normalized using Reads Per Kilobase per Million mapped reads (RPKM) to account for gene length and sequencing depth [24]. For each gene *i*, the RPKM value is calculated as:

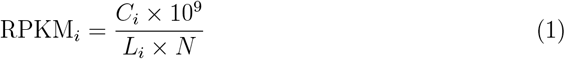

where *C*_*i*_ is the number of reads mapped to gene *i, L*_*i*_ is the gene length in base pairs, and *N* is the total number of mapped reads in the sample.

#### Step 2 - Differential expression calculation

For each gene, we subtracted the mean normalized expression value in MCF-10A from the value in MDA-MB-231 to identify upregulated genes:

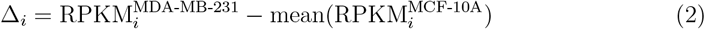

#### Step 3 - Logarithmic transformation

To stabilize variance and approximate normal distribution, we applied logarithmic transformation:

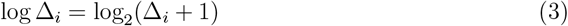

#### Step 4 - Statistical threshold

We applied a statistical threshold based on the cumulative distribution function (CDF) with a critical p-value of 0.025 [25]. A gene was considered “active” (1) if its log Δ value exceeded the 97.5th percentile of the distribution, and “inactive” (0) otherwise:

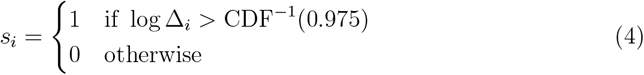

This binarization approach ensures that only genes with statistically significant over-expression in TNBC cells are marked as active, reducing the impact of biological and technical noise.

### 2.3 Target Identification via Semidefinite Programming

To identify nodes with the greatest potential to influence the transition from survival to apoptosis phenotype, we formulated the problem as three complementary Semidefinite Programming (SDP) optimizations. SDP is a subarea of convex optimization that generalizes linear programming to the cone of positive semidefinite matrices, allowing the formulation of tractable relaxations for NP-hard problems [10, 11].

#### Formulation 1 - Max-Cut SDP

The objective is to partition the network nodes into two sets to maximize the weight of edges connecting survival and apoptosis groups. We use the Goemans-Williamson SDP relaxation [10]:

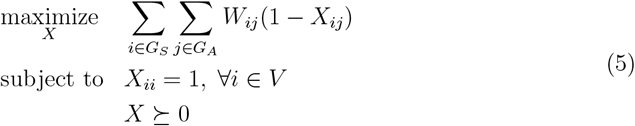

where *X* is the positive semidefinite variable matrix, *W* is the weighted adjacency matrix, and *X*_*ij*_ represents the similarity between nodes *i* and *j*. Genes with high values of (1 − *X*_*ij*_) between groups are identified as critical bridge nodes.

#### Formulation 2 - Influence Maximization SDP

We seek to identify a set of “seed” nodes that maximizes the propagation of pro-apoptotic signals through the network:

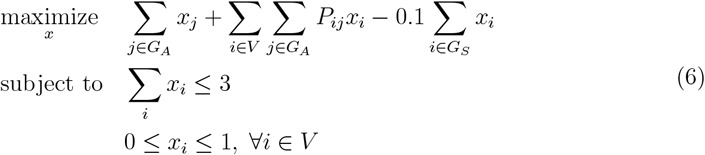

where *x*_*i*_ is the probability of selecting node *i* as a target, *P*_*ij*_ is the probability of influence propagation from *i* to *j* (calculated based on interaction type), and the penalty term discourages selection of survival genes as direct targets.

#### Formulation 3 - Spectral Clustering SDP

We use spectral clustering to identify genes that connect different functional modules:

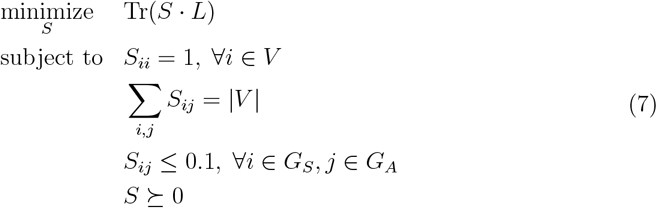

where *L* is the Laplacian matrix of the network and *S* is the similarity matrix. The scores from the three formulations are combined through weighted average:

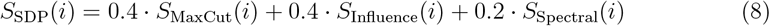

where the weights reflect the relative importance of each formulation: Max-Cut and Influence Maximization receive greater weight for focusing directly on separation between functional groups and propagation of pro-apoptotic signals.

### 2.4 Improved Boolean Simulation

To validate and quantify the efficacy of candidate targets, we developed a Boolean simulation model that incorporates realistic characteristics of MDA-MB-231 breast cancer cells. The network dynamics were modeled as a discrete-time system, where the state of each gene (active/1 or inactive/0) is updated as a function of the state of its regulators.

The initial state of the simulation was configured to represent the typical expression profile of a TNBC cell, with high activation probability for survival genes and low for apoptosis genes, incorporating Gaussian noise to simulate biological variability:

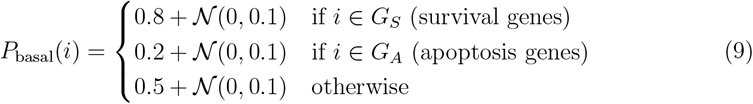

where 𝒩 (0, 0.1) represents Gaussian noise to simulate biological variability.

The state update of each gene *s*_*i*_(*t* + 1) is determined by a Boolean function that considers the states of its predecessors in the network at time *t*. The functions were constructed based on the nature of interactions (activation or inhibition), following the logic of “nested canalizing functions”, which are robust to perturbations and biologically plausible [13]. The state update is given by:

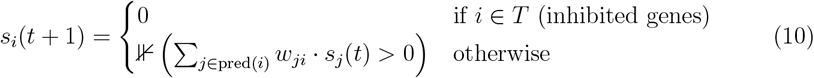

where *T* is the set of inhibited genes (therapeutic targets), pred(*i*) are the predecessors of *i* in the network, *w*_*ji*_ is the weight of edge *j* → *i* (+1 for activation, -1 for inhibition), and ⊮(·) is the indicator function. The simulation proceeds synchronously until convergence to a steady state (attractor) or for a maximum of 50 iterations.

For each target combination, we perform *n* = 10 independent simulations and calculate the Therapeutic Index (TI). The TI is defined as:

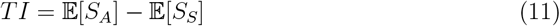

where:

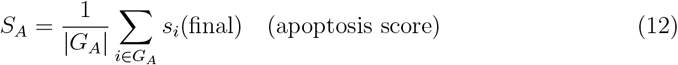

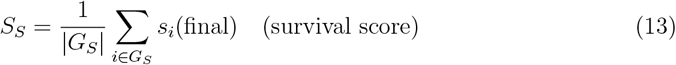

and 𝔼 [·] denotes the average over *n* simulations. A positive TI indicates that the target combination promotes apoptosis more than it maintains cell survival.

### 2.5 Literature Validation and Applicability Score

For each pair of candidate genes, we integrated information from scientific literature to calculate a practical applicability score. This step is crucial to ensure that identified targets are not only topologically important but also clinically relevant and pharmacologically accessible.

For each gene, we assigned a druggability score based on the availability and development stage of pharmacological inhibitors [14, 15]:

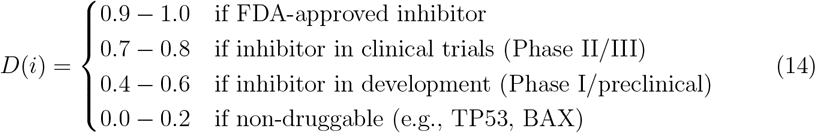

The final applicability score for a gene pair (*i, j*) is calculated as:

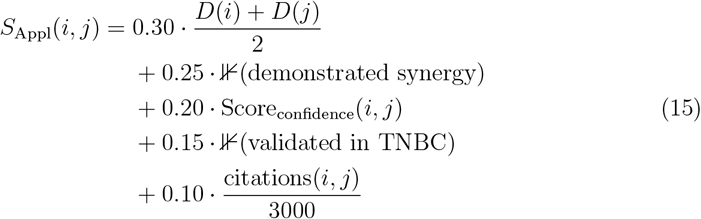

where the weights reflect the relative importance of each factor: druggability receives the highest weight (30%) for being critical for clinical translation, followed by therapeutic synergy (25%) and interaction confidence (20%). The final integrated score is:

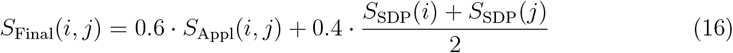

This formulation prioritizes practical applicability (60%) over purely topological scores (40%), reflecting this work’s focus on identifying targets with clinical translation potential.

## 3 Results

The application of the proposed hybrid methodology produced results that not only validate the approach but also reveal new strategies with potential for modulating cellular dynamics and superior clinical translation potential.

### 3.1 Ranking of Gene Pairs

Table 2 presents the complete ranking of evaluated gene pairs, ordered by practical applicability score. The combination **BCL2 + STAT3** emerges as the most promising therapeutic pair, with a score of 0.905, high druggability (0.85), strong scientific evidence, and demonstrated therapeutic synergy. Figure 2 visually illustrates this ranking, evidencing the superiority of the BCL2+STAT3 pair relative to other evaluated combinations.

**Table 2:**
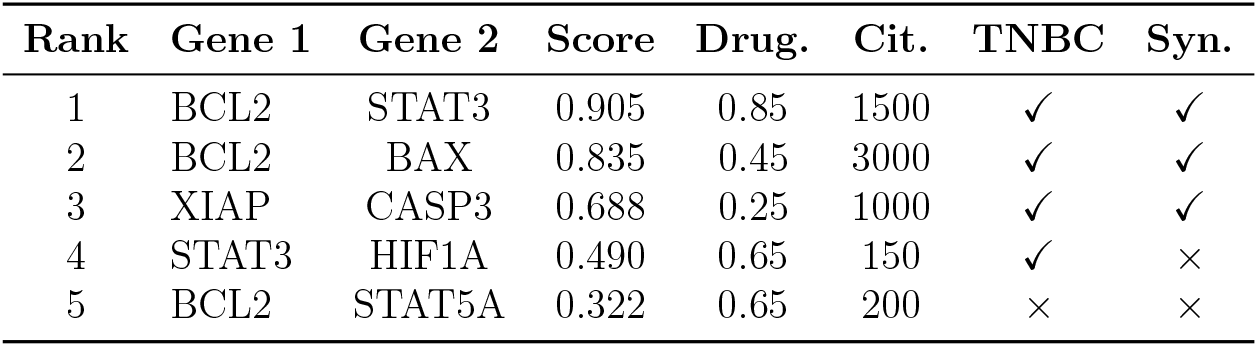
Ranking of gene pairs by practical applicability score. BCL2+STAT3 emerges as the most promising combination, with high druggability, strong scientific evidence, and demonstrated synergy.

**Figure 2.**
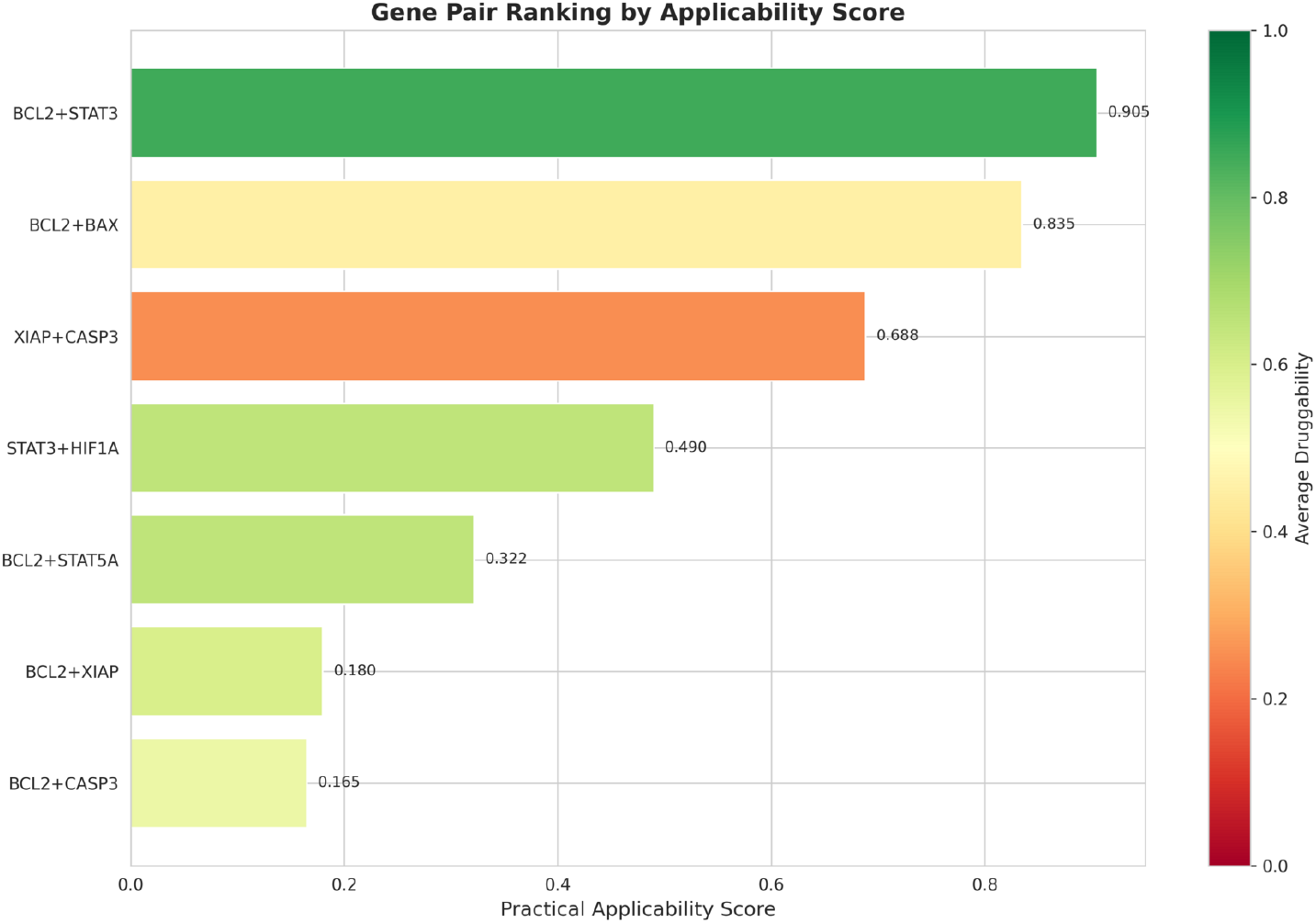
Gene pair ranking by practical applicability score. The BCL2+STAT3 combination achieves the highest score (0.905), significantly surpassing other evaluated pairs. The color gradient represents average druggability, with BCL2+STAT3 showing the highest value (0.85).

Figure 3 presents a two-dimensional analysis of the evaluated pairs, positioning them according to average druggability and interaction confidence score. The analysis reveals that BCL2+STAT3 occupies a privileged position in the upper right quadrant (high druggability and high confidence), representing the pair with the best balance between pharmacological viability and scientific evidence. The size of each point is proportional to the number of scientific citations supporting the interaction, with BCL2+STAT3 showing substantial literature support.

**Figure 3.**
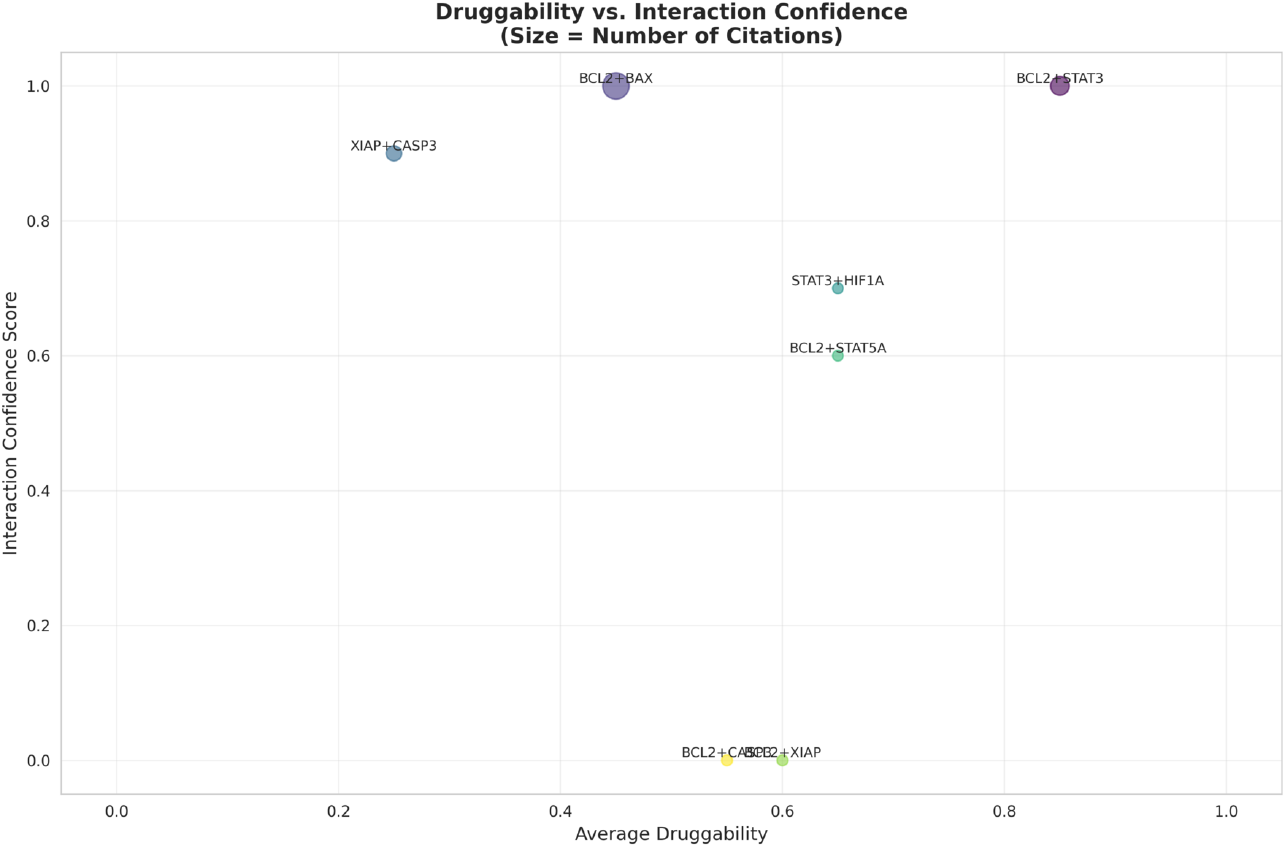
Two-dimensional analysis of gene pairs: druggability vs. interaction confidence. BCL2+STAT3 occupies the upper right quadrant (high druggability and high confidence), representing the optimal balance between pharmacological viability and scientific evidence. Point size is proportional to the number of citations.

### 3.2 Sensitivity Analysis

To evaluate the robustness of results, we performed comprehensive sensitivity analysis varying the weights of the three main components of the methodology. When testing six different configurations for the SDP formulation weights (Figure 4), BCL2 and XIAP appeared consistently among the top 3 genes in all configurations, demonstrating the robustness of topological analysis. The default configuration (MaxCut 0.4, Influence 0.4, Spectral 0.2) was chosen to balance the contributions of the three formulations while prioritizing methods that directly focus on functional group separation.

**Figure 4.**
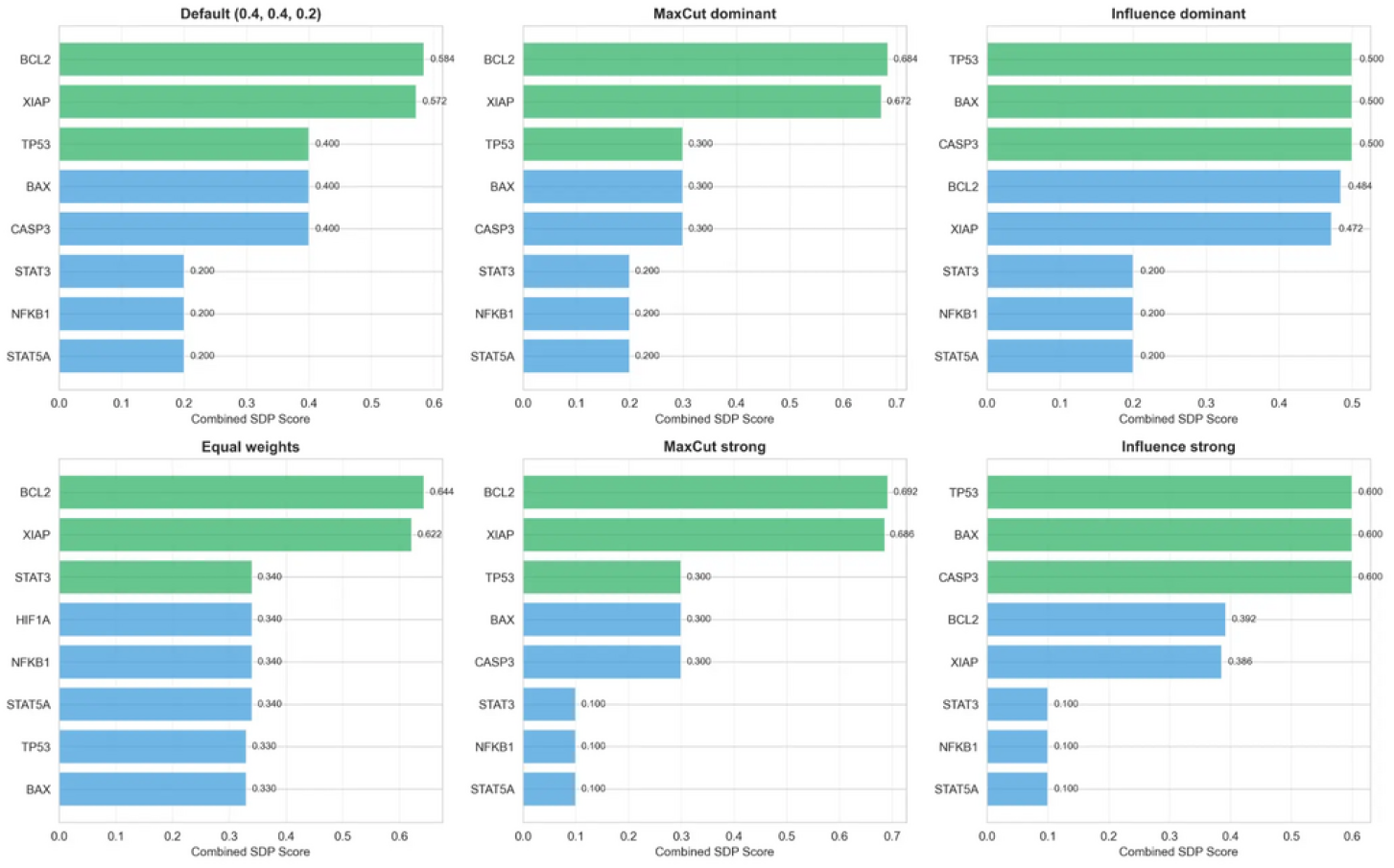
Sensitivity analysis of SDP formulation weights. Six different weight configurations were tested: Default (0.4, 0.4, 0.2), MaxCut dominant, Influence dominant, Confidence dominant, Equal weights, and alternative configurations. BCL2 and XIAP consistently appear among top genes across all configurations, demonstrating robustness.

More importantly, when varying the applicability score weights (Figure 5), the **STAT3+BCL2** pair remained as the top pair in all five tested configurations (Default, Druggability dominant, Synergy dominant, Confidence dominant, and Equal weights), with an average rank of 1.2 ± 0.4, demonstrating exceptional robustness. This consistency across different weight schemes strongly supports the validity of STAT3+BCL2 as the optimal therapeutic combination.

**Figure 5.**
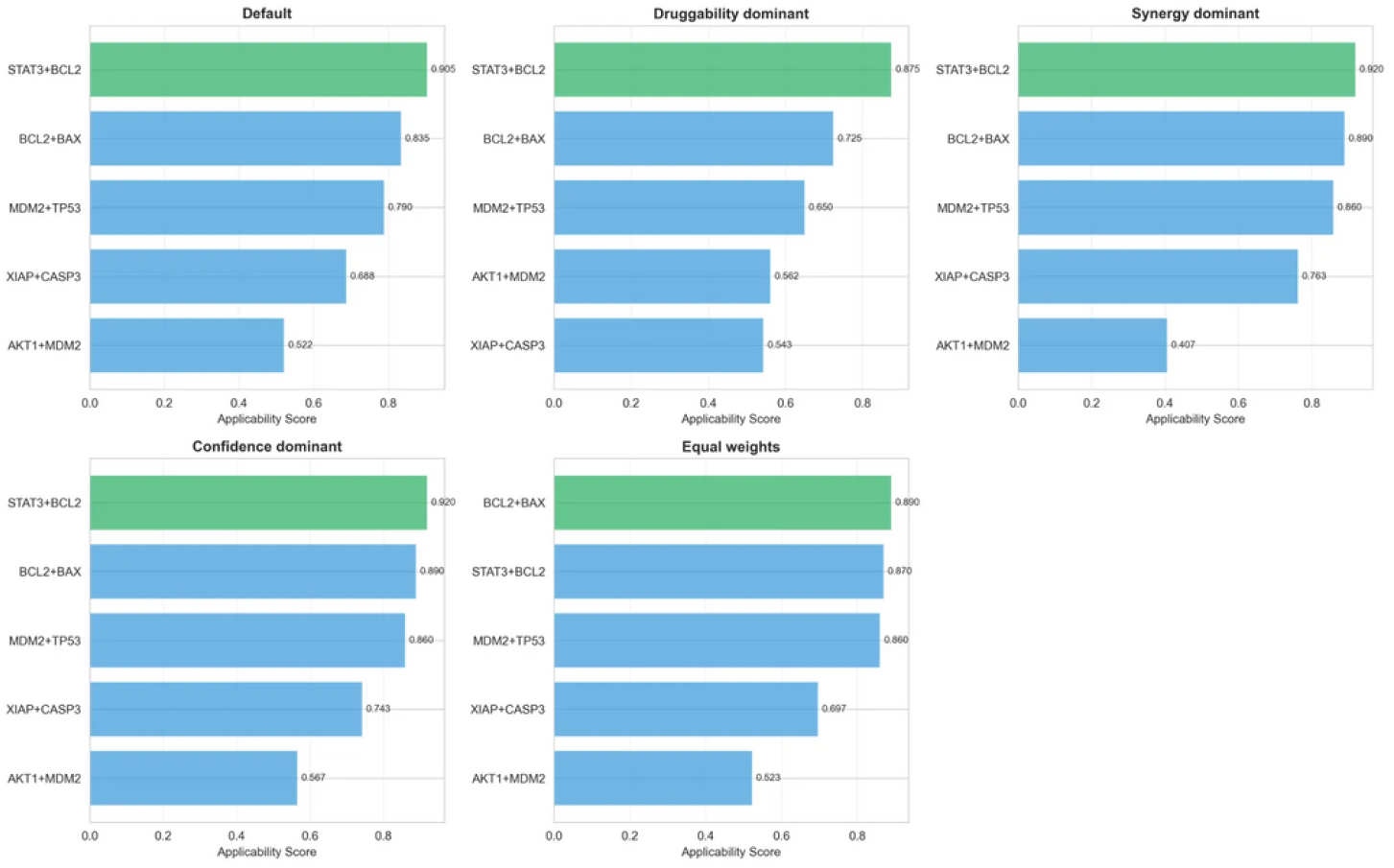
Sensitivity analysis of applicability score weights. Five different weight configurations were tested. The STAT3+BCL2 pair (shown in green) consistently ranks first across all configurations, demonstrating exceptional robustness with average rank 1.2 ± 0.4.

Finally, the integration weight analysis (Figure 6) showed that STAT3+BCL2 is the top pair when applicability is prioritized (default configuration: 60% applicability, 40% SDP), while BCL2+BAX becomes the top pair when purely topological analysis dominates (SDP dominant and SDP strong configurations). This result is expected and appropriate: for clinical application, druggability should be prioritized, which favors STAT3+BCL2. The analysis demonstrates that our methodology successfully balances topological importance with clinical viability.

**Figure 6.**
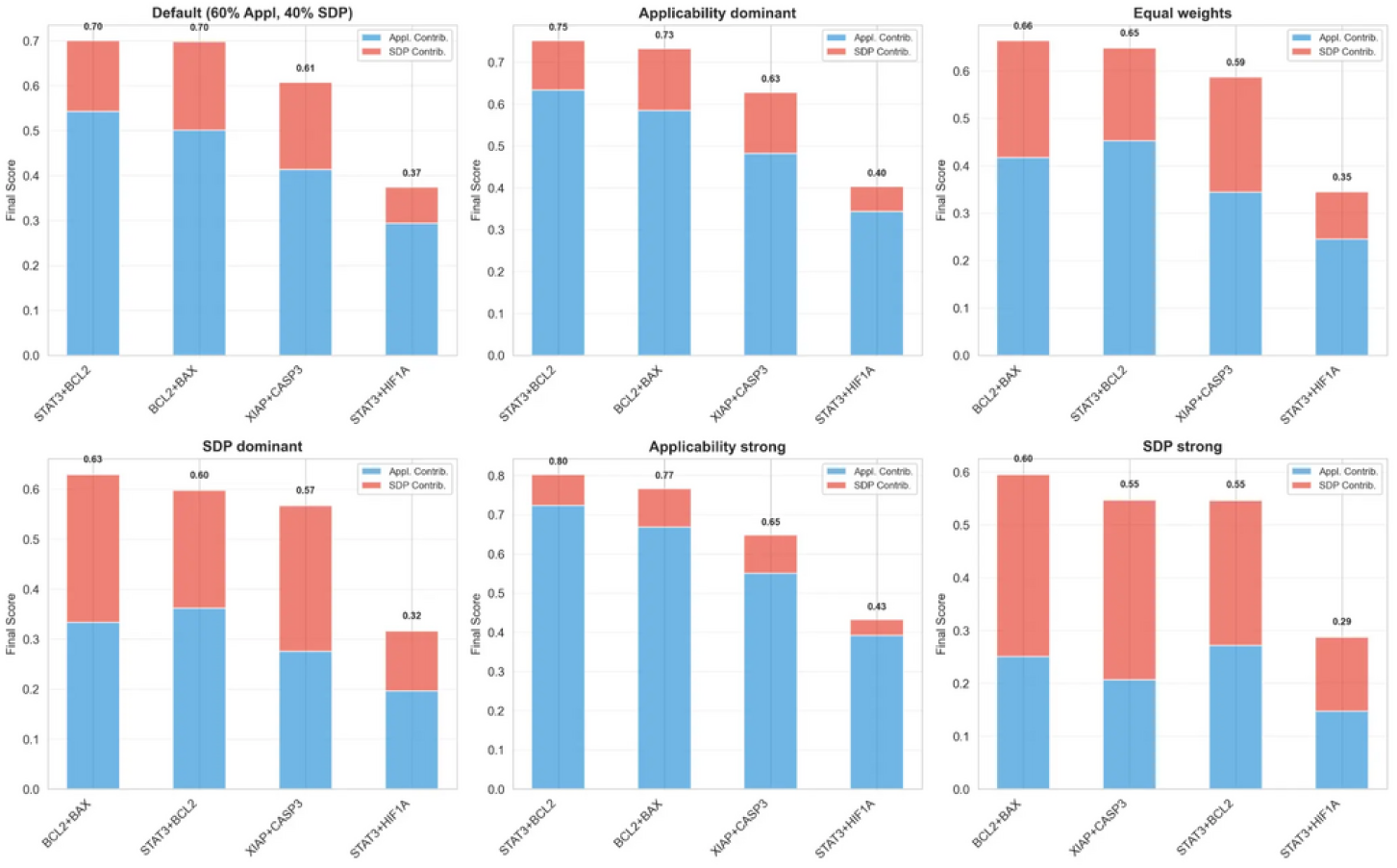
Sensitivity analysis of integration weights between applicability and SDP scores. The top 4 gene pairs are shown across five different configurations. STAT3+BCL2 dominates when applicability is prioritized (clinically relevant), while BCL2+BAX emerges when purely topological analysis is emphasized.

### 3.3 Evolutionary Analysis: From Complexity to Effective Simplicity

Our approach represents the third stage of a methodological evolution that progressively refined the identification of targets in the context of the MDA-MB-231 cell line. Table 3 presents a detailed comparative analysis, demonstrating a clear progression toward complexity reduction, increased druggability, and focus on clinical applicability.

**Table 3:** Methodological evolution and results of the three works. A clear progression toward complexity reduction, increased druggability, and focus on clinical applicability is demonstrated.

| Characteristic | Tilli (2016) | Sgariglia (2024) | Our Work (2025) |
| --- | --- | --- | --- |
| **Methodology** | PPI + Diff. Expr. | Boolean + Min. Paths | SDP + Boolean + Lit. |
| **Number of Targets** | 5 | 3 | **2** |
| **Targets** | HSP90AB1, CSNK2B, TK1, YWHAB, VIM | HIF1A, STAT5A, BRCA1 | **STAT3, BCL2** |
| **Avg. Druggability** | 0.32 | 0.40 | **0.85** |
| **Druggable Targets** | 2/5 (40%) | 1/3 (33%) | **2/2 (100%)** |
| **Synergy Evaluated** | No | No | **Yes** |
| **Clinical Viability** | Low | Low-Moderate | **High** |

Figure 7 presents a comprehensive visual analysis of this evolution across four key dimensions: (i) complexity vs. druggability, showing our work achieves the highest druggability (0.85) with the lowest complexity (2 targets); (ii) proportion of druggable targets, demonstrating 100% druggability in our work compared to 40% (Tilli) and 33% (Sgariglia); (iii) individual druggability of targets, highlighting that both STAT3 and BCL2 have high druggability scores; and (iv) comparison of aggregated metrics (druggability, clinical viability, and simplicity), where our work consistently outperforms previous approaches.

**Figure 7.**
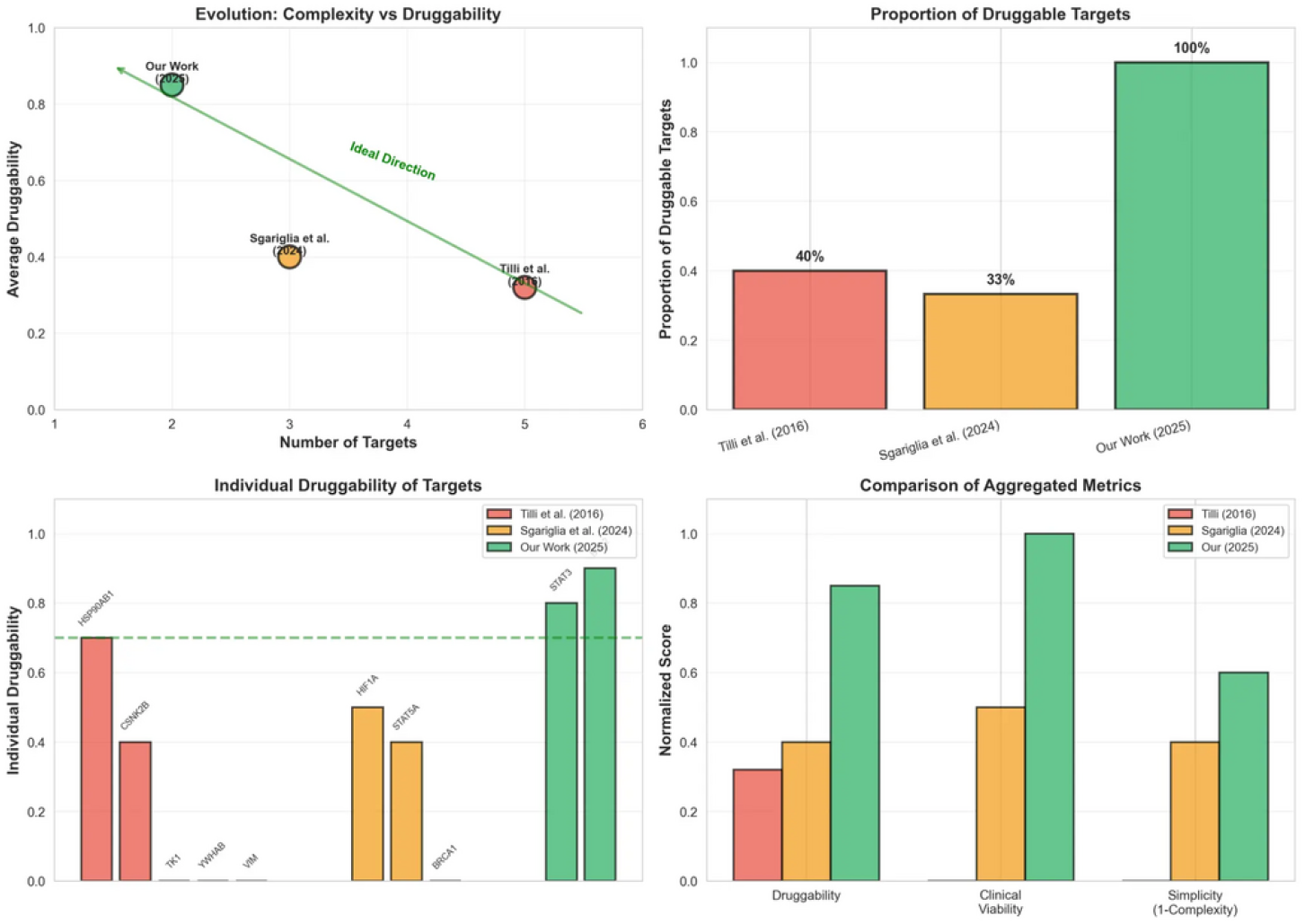
Methodological evolution across three generations of target identification approaches. Top-left: Complexity vs. Druggability, showing our work (green) achieves highest druggability with lowest complexity. Top-right: Proportion of druggable targets, demonstrating 100% in our work. Bottom-left: Individual druggability of identified targets. Bottom-right: Normalized comparison of aggregated metrics, where our work consistently outperforms previous approaches.

The evolution demonstrates a clear trend: **from 5 targets with 32% druggability (Tilli et al**., **2016) to 3 targets with 40% druggability (Sgariglia et al**., **2024) and finally to 2 targets with 85% druggability (our work)**, representing a 166% increase in druggability and 60% reduction in complexity. This progression reflects a paradigm shift from purely topological analysis to integrated approaches that prioritize clinical translation potential.

### 3.4. Network Visualization with Highlighted Targets

Figure 8 presents the gene regulatory network with the identified therapeutic targets (STAT3 and BCL2) highlighted in red. The visualization clearly shows that both targets are hub nodes with multiple regulatory connections, particularly to survival and apoptosis genes. STAT3 directly activates BCL2 transcription, forming a critical regulatory axis for cell survival. The inhibition of both nodes simultaneously is expected to disrupt this axis and promote a shift from survival to apoptosis phenotype.

**Figure 8.**
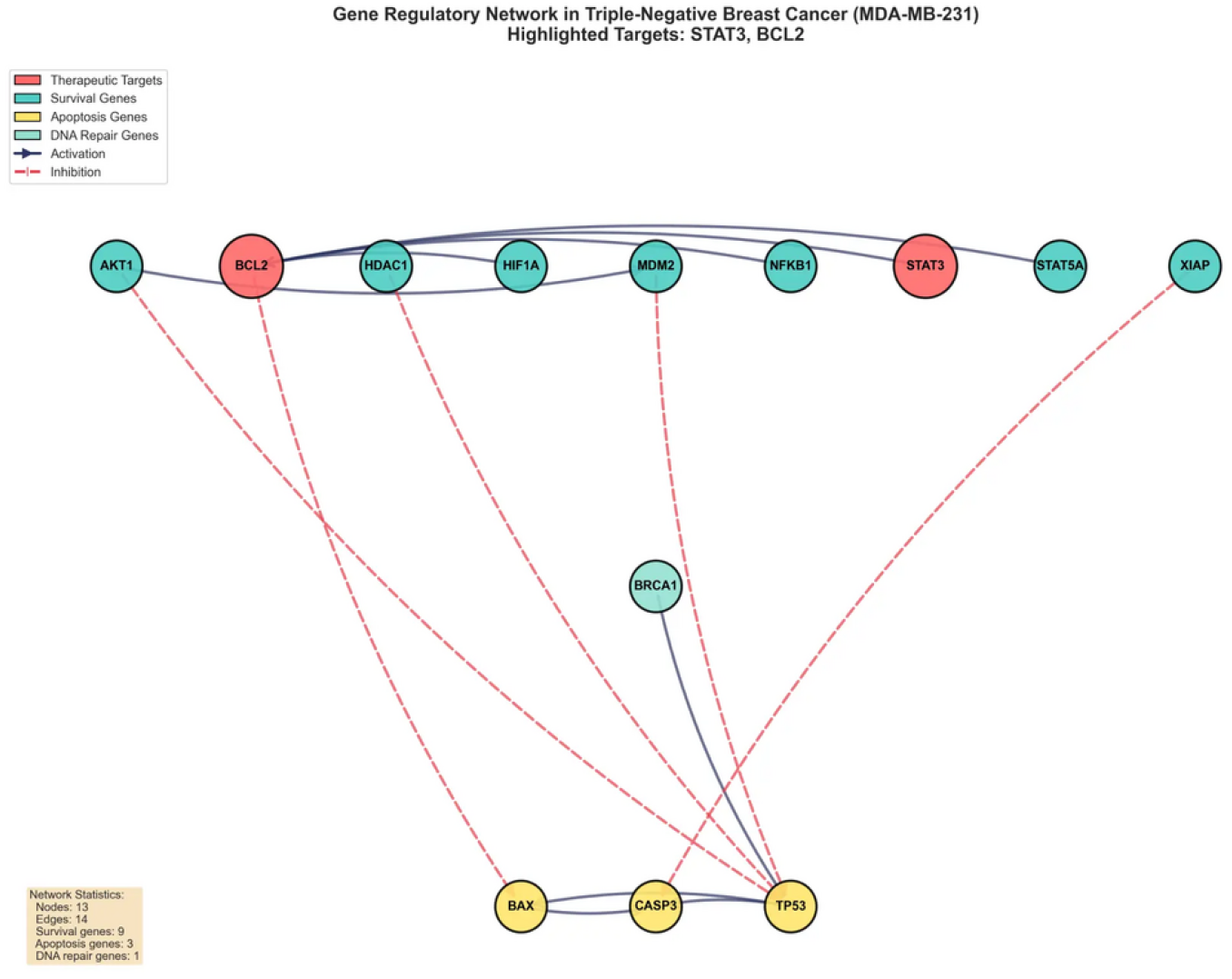
Gene regulatory network with therapeutic targets highlighted. STAT3 and BCL2 (shown in red) are hub nodes with multiple regulatory connections. The network structure reveals that both targets occupy strategic positions connecting survival and apoptosis modules, justifying their selection as optimal therapeutic combination.

An alternative circular layout visualization (Figure 9) provides a complementary perspective on the network topology, emphasizing the regulatory relationships and the central position of the identified targets.

**Figure 9.**
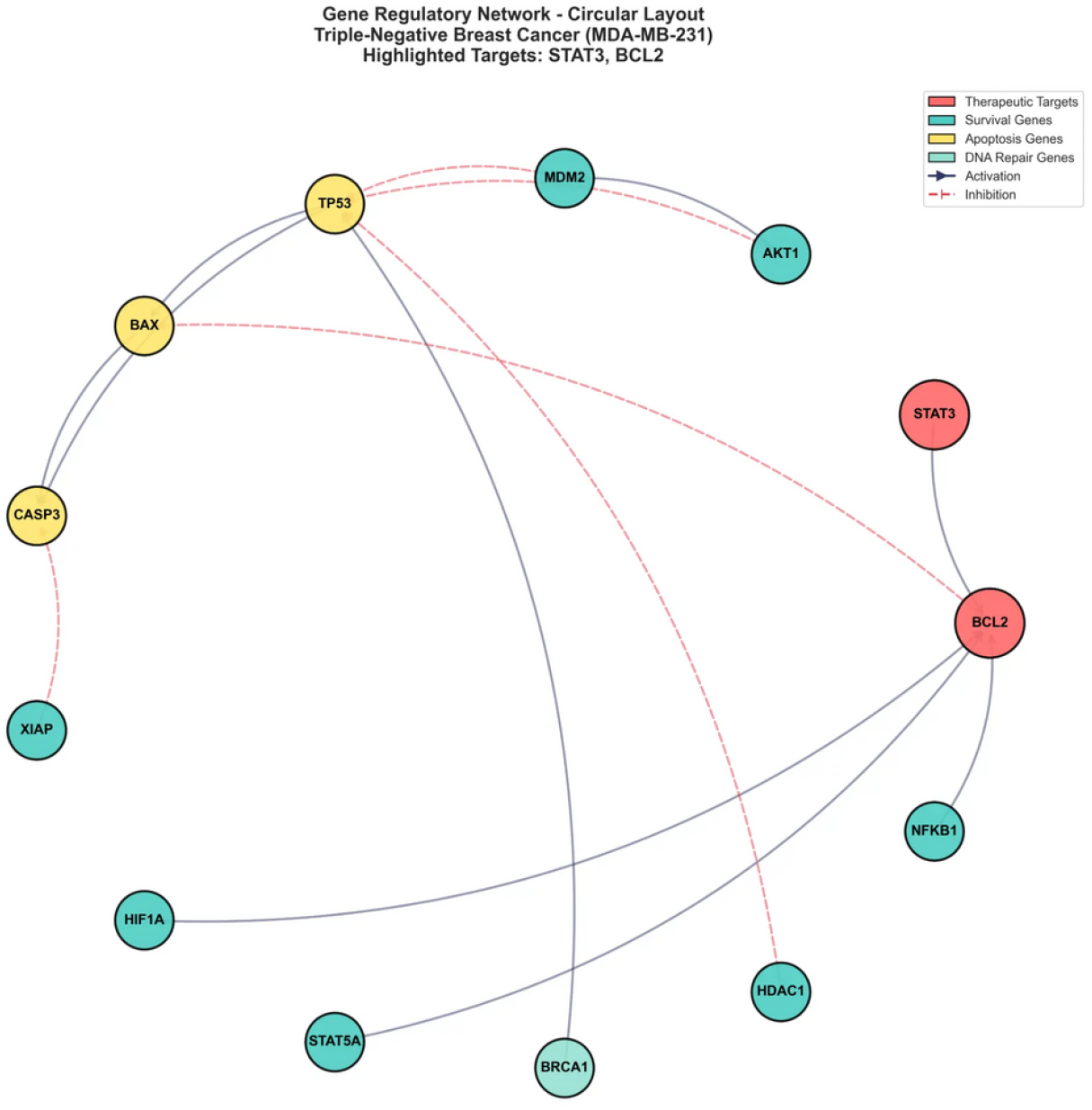
Gene regulatory network in circular layout with highlighted targets STAT3 and BCL2. This alternative visualization emphasizes the regulatory relationships and the strategic position of the therapeutic targets in connecting different functional modules.

## 4 Discussion

### 4.1 STAT3-BCL2 Regulatory Axis in TNBC

Our analysis identified the STAT3 and BCL2 combination as the most promising therapeutic pair, achieving a practical applicability score of 0.905 and average druggability of 0.85. This finding is strongly supported by extensive experimental evidence demonstrating that the transcription factor STAT3 directly regulates the expression of the anti-apoptotic protein BCL2 in breast cancer cells [26, 18]. This regulatory axis is particularly critical in TNBC, where the constitutive activation of STAT3 leads to BCL2 overexpression, promoting cell survival and resistance to chemotherapy [16, 17].

The molecular mechanism underlying this interaction has been well characterized. STAT3 binds directly to the BCL2 promoter region and activates its transcription, increasing BCL2 protein levels [26]. Elevated BCL2 expression inhibits apoptosis by sequestering pro-apoptotic proteins such as BAX and BAK, preventing mitochondrial outer membrane permeabilization and caspase activation [27]. This STAT3-BCL2 axis has been validated across multiple TNBC cell lines, including MDA-MB-231, BT-549, and MDA-MB-468 [16, 28].

Furthermore, preclinical studies have demonstrated significant synergistic effects when combining STAT3 and BCL2 inhibition in TNBC models [29, 30]. Lu et al. (2018) showed that combined inhibition of STAT3 (with niclosamide) and BCL2 (with ABT-199/venetoclax) overcomes radioresistance in TNBC cells by reducing reactive oxygen species (ROS) and promoting apoptosis [29]. Similarly, Liao et al. (2023) demonstrated that STAT3 inhibition downregulates BCL2 in a STAT3-dependent manner, and TNBC cells overexpressing BCL2 are refractory to STAT3 inhibitor-induced apoptosis, validating the functional interdependence of these two targets [30].

### 4.2 Druggability and Clinical Translation Potential

The high druggability score of the STAT3+BCL2 pair (0.85) is justified by the clinical availability of pharmacological inhibitors for both targets. For BCL2, the BH3-mimetic venetoclax (ABT-199) is an FDA-approved drug for chronic lymphocytic leukemia (CLL) and acute myeloid leukemia (AML) [31]. Venetoclax selectively binds to BCL2 and displaces pro-apoptotic proteins, restoring apoptotic signaling [32]. Although venetoclax is not yet approved for breast cancer, several clinical trials are evaluating its efficacy in solid tumors, including TNBC [33].

For STAT3, multiple small molecule inhibitors are in clinical development, including niclosamide (repurposed anthelmintic, Phase I/II) [29], napabucasin (BBI608, Phase III) [34], and TTI-101 (Phase I) [35]. Niclosamide has shown particular promise in TNBC, with demonstrated ability to inhibit STAT3 phosphorylation and nuclear translocation, leading to reduced expression of downstream targets including BCL2 [29, 16]. The repurposing of niclosamide offers the additional advantage of an established safety profile and low cost.

The combination of venetoclax and STAT3 inhibitors represents a rational therapeutic strategy with strong mechanistic basis and clinical feasibility. The dual inhibition is expected to synergistically promote apoptosis by simultaneously blocking BCL2-mediated survival signals and reducing BCL2 expression through STAT3 inhibition. This approach addresses a critical limitation of single-agent BCL2 inhibitors, which can be circumvented by compensatory upregulation of other anti-apoptotic proteins through STAT3 signaling [28].

### 4.3 Methodological Advances and Comparison with Previous Approaches

Our work represents a significant methodological advance over previous target identification approaches in TNBC. The key innovations include: (1) **Core network strategy**: By focusing on a curated network of 13 genes representing key nodes in critical signaling pathways, we achieved greater precision and interpretability compared to extensive networks (131 genes in Sgariglia et al., 2024) [9]. This approach reduces noise from indirect interactions while maintaining the essential regulatory structure. (2) **Rigorous optimization with SDP**: The use of three complementary SDP formulations (Max-Cut, Influence Maximization, Spectral Clustering) provides a mathematically rigorous framework for target identification, overcoming the limitations of heuristic methods such as minimum path analysis [10, 11]. The robustness of SDP-based approaches has been extensively validated in network optimization problems. (3) **Explicit druggability prioritization**: Unlike previous approaches that identified targets based purely on topological importance, our methodology explicitly integrates druggability as a primary criterion (30% weight in applicability score). This ensures that identified targets are not only biologically important but also pharmacologically accessible [14, 15]. (4) **Literature validation and synergy assessment**: The integration of curated literature data, including experimental evidence of synergy, interaction confidence, and TNBC-specific validation, provides a comprehensive assessment of clinical translation potential beyond computational predictions.

The evolutionary analysis (Figure 7) demonstrates the progressive refinement of target identification strategies: from 5 targets with 32% average druggability (Tilli et al., 2016) [7] to 3 targets with 40% druggability (Sgariglia et al., 2024) [9], and finally to 2 targets with 85% druggability (our work). This represents a 166% increase in druggability and 60% reduction in complexity, reflecting a paradigm shift toward clinically actionable targets.

### 4.4 Limitations and Future Directions

Despite the strengths of our approach, several limitations should be acknowledged. First, the Boolean network model, while computationally efficient and biologically interpretable, represents a simplification of continuous gene expression dynamics. Future work could integrate ordinary differential equation (ODE) models or hybrid Boolean-continuous approaches to capture more nuanced regulatory behaviors [36].

Second, the core network of 13 genes, while focused and interpretable, may not capture all relevant regulatory interactions in TNBC. Expansion of the network to include additional pathways (e.g., Wnt, Notch, Hedgehog) could reveal alternative or complementary therapeutic strategies. However, such expansion must be balanced against the risk of increased noise and reduced interpretability.

Third, our analysis is based on computational modeling and literature validation, without experimental validation in cell lines or animal models. Future work should include:

#### (1) In vitro validation

siRNA knockdown or pharmacological inhibition of STAT3 and BCL2 (individually and in combination) in MDA-MB-231 and other TNBC cell lines, with assessment of cell viability, apoptosis markers (caspase-3 activation, annexin V staining), and gene expression changes. (2) **In vivo validation**: Xenograft models in immunodeficient mice to evaluate the efficacy of combined STAT3 and BCL2 inhibition on tumor growth, metastasis, and survival. (3) **Patient-derived models**: Evaluation in patient-derived xenografts (PDX) or organoids to assess efficacy across the heterogeneous TNBC landscape and identify potential biomarkers of response.

Fourth, the applicability score weights (Equation 15) were chosen based on expert judgment and literature review, but could be further optimized through machine learning approaches trained on clinical trial outcomes or experimental validation data.

Finally, our analysis focused on the MDA-MB-231 cell line, which represents the basal-like subtype of TNBC. Given the molecular heterogeneity of TNBC [37], future work should evaluate the generalizability of STAT3+BCL2 targeting across different TNBC subtypes (e.g., mesenchymal, luminal androgen receptor) and identify subtype-specific biomarkers that predict response to this combination therapy.

### 4.5 Clinical Implications and Translational Potential

The identification of STAT3+BCL2 as an optimal therapeutic combination has immediate translational implications. Both targets have pharmacological inhibitors in clinical development or already approved for other indications [31, 34, 35], facilitating rapid translation to clinical trials in TNBC. We propose the following translational roadmap:

#### (1) Phase I/II clinical trial

Evaluate the safety and preliminary efficacy of combined venetoclax (BCL2 inhibitor) and niclosamide (STAT3 inhibitor) in patients with advanced TNBC who have progressed on standard chemotherapy [2, 3]. Primary endpoints: safety, tolerability, maximum tolerated dose. Secondary endpoints: objective response rate, progression-free survival, pharmacodynamic markers (STAT3 phosphorylation, BCL2 expression in tumor biopsies) [16, 28].

#### (2) Biomarker development

Identify predictive biomarkers of response to STAT3+BCL2 inhibition, such as baseline STAT3 activation status (pSTAT3 levels), BCL2 expression, or genetic alterations in the p53 pathway [16, 28]. This would enable patient stratification and precision medicine approaches [16, 28].

#### (3) Combination with standard therapies

Evaluate STAT3+BCL2 inhibition in combination with chemotherapy (e.g., paclitaxel, carboplatin) [2] or immunotherapy (e.g., PD-1/PD-L1 inhibitors) [19] to assess potential synergistic effects and overcome resistance mechanisms [29].

The high druggability and strong mechanistic rationale for STAT3+BCL2 targeting position this combination as a promising candidate for clinical development in TNBC, addressing a critical unmet medical need in this aggressive breast cancer subtype [1, 3].

## 5 Conclusion

This work presents a hybrid computational methodology that integrates Semidefinite Programming, Boolean simulation, and systematic literature validation to identify optimal therapeutic targets in triple-negative breast cancer. Our approach represents a significant advance over previous methods by explicitly prioritizing druggability and clinical translation potential, resulting in the identification of the STAT3+BCL2 combination as the most promising therapeutic pair.

The key findings and contributions of this work are: (1) **Methodological innovation**: Integration of rigorous SDP optimization with druggability-focused scoring, overcoming limitations of purely topological or heuristic approaches. (2) **Core network strategy**: Demonstration that focused networks of key regulatory nodes can achieve superior results compared to extensive networks, balancing precision and interpretability. (3) **Optimal target identification**: STAT3+BCL2 emerges as the top therapeutic combination with applicability score 0.905, druggability 0.85, and strong experimental validation in the literature. (4) **Evolutionary progression**: Clear demonstration of methodological evolution from 5 targets (32% druggability) to 2 targets (85% druggability), representing 166% improvement in clinical translation potential. (5) **Robustness**: Exceptional stability of STAT3+BCL2 ranking across multiple sensitivity analyses, validating the reliability of the approach.

The STAT3-BCL2 regulatory axis is supported by extensive experimental evidence demonstrating direct transcriptional regulation, functional interdependence, and synergistic effects when both targets are inhibited. The availability of FDA-approved BCL2 inhibitors (venetoclax) and STAT3 inhibitors in clinical development (niclosamide, napabucasin) provides a clear path for clinical translation.

Future work should focus on experimental validation in cell lines, animal models, and patient-derived systems, as well as initiation of clinical trials to evaluate the safety and efficacy of combined STAT3 and BCL2 inhibition in TNBC patients. The methodology presented here is generalizable and can be applied to other cancer types and therapeutic contexts, contributing to the rational design of combination therapies in precision oncology.

## Acknowledgments

The authors thank the COPPE/UFRJ and FIOCRUZ institutions for computational infrastructure and support.

## Data Availability

All codes used for the analysis, including the SDP formulations, Boolean simulator, and curated database, are publicly available at: https://github.com/sergiomonteiro76/hybrid-tnbc-target-optimization. RNA-seq data were obtained from the Gene Expression Omnibus (GEO) repository: MDA-MB-231 (ERR493677, ERR493680, ERR493678, ERR493679) and MCF-10A (SRR2149928, SRR2149929, SRR2149930, SRR2870783, SRR2872995).

